# Glucose derived redox equivalents preserve PKA activity and glucagon secretion during hypoglycaemia

**DOI:** 10.64898/2026.08.11.744097

**Authors:** Alexander Frueh, Georgios Katzilieris-Petras, Caroline L. Pedersen, Maia H. Ekstrand, Ganga Deshar, Renata Ialchina, Hayden A Paige, Dorthe Nielsen, Daniel B. Andersen, Jens J. Holst, Peter Spegel, Per A. Pedersen, Jakob G. Knudsen

## Abstract

The release of glucagon from pancreatic alpha cells is a core component of hypoglycaemic counter regulation. Several mechanisms regulate glucagon release including paracrine control by neighbouring cell types, and changes in extracellular glucose. While the inhibitory effect of glucose on glucagon secretion is well established, the exact way in which glucose metabolism contributes to alpha cell function remains unclear. Here, we use live-cell imaging of the redox potential in alpha cells within intact islets to investigate whether non-mitochondrial glucose metabolism contributes to the potentiation of glucagon secretion at low glucose. Our findings show that increased glucose metabolism through the pentose phosphate pathway elevates the cytosolic redox potential in alpha cells. Using a combination of antioxidant treatment and pre-incubation in 5 mM glucose, we find that the cytosolic redox potential affects PKA activity in alpha cells and that changes in whole body redox state affects the counterregulatory response in mice. These findings indicate that prior glucose-driven redox potential charging is essential for maintaining glucagon secretion at low glucose.

## Introduction

Pancreatic alpha cells release glucagon in response to low blood glucose to promote catabolic processes in multiple tissues, with the most prominent effect occurring in the liver to support hepatic glucose production (1–4). However, in individuals with diabetes, glucagon secretion becomes dysregulated, with alpha cells releasing higher amounts of glucagon under normoglycaemia (5–7), while counterregulatory glucagon responses become inadequate (8).

The initiation of glucagon secretion from alpha cells during hypoglycaemia is thought to rely mainly on two mechanisms; an electrically induced activation of exocytosis (9) and paracrine or endocrine amplification of granule release through intracellular signalling (10–12). While the change in electrical activity is thought to be regulated through the metabolic action of glucose (13–16), alpha cells exhibit limited glucose oxidation (17; 18) and instead rely on fatty acid oxidation to provide ATP (14; 19-21). Previous observations suggest that total glucose utilisation is similar in beta cells and non-beta cells (17), implying that glucose uptake and non-oxidative glucose metabolism may play an important role in alpha cell function, despite limited glucose oxidation. Here, we explored the role of non-oxidative glucose metabolism in the regulation of glucagon secretion.

## Materials and Methods

### Animals

All animal experiments were approved by the Danish Animal Inspectorate. Female C57BL6/Njr mice (Janvier) were housed on a 12–12-h light-dark cycle at 22°C with access to ad libitum standard chow and water. Mice were used for experiments at 12-20 weeks of age.

### Islet isolation

Mice were euthanized by cervical dislocation and 0.09 mg/ml of ice cold Liberase (Roche, 05401020001) in Hanks buffered saline solution (Sigma Aldrich, H6648) was injected into the bile duct after clamping the pancreatic duct at the Ampullae of Vater. Pancreata were then isolated and kept on ice before digestion in a 37°C water bath for 12 minutes. Ice cold Hank’s solution supplemented with 0.2% BSA was used to stop the digestion, and islets were mechanically dissociated from exocrine tissue by shaking. Islets were handpicked into RPMI media (Gibco, 11879-020) supplemented with 7 mM glucose, 10% FBS (Sigma Aldrich, F7524) and 1% penicillin/streptomycin (P/S; Gibco 15140-122) and cultured at 5% CO_2_, 37°C for 1h before initiation of experiments. All subsequent experiments were performed in Krebs Ringer Buffer (KRB; 115 mmol/L NaCl, 4.6 mmol/L KCl, 2.6 mmol/L CaCl2, 1.2 mmol/L MgCl2, 1 mmol/L NaH2PO4, 25 mmol/L NaHCO3, 10 mmol/L HEPES, pH7.4, 6.6 mg/mL FA-free BSA and 0.36 mM non esterified fatty acids) according to (14).

### Metabolomics

Metabolomics on alphaTC1-6 cells were performed as previously described [22]

### Generation of viral constructs

To generate viral constructs for alpha cell specific expression of life cell imaging probes, sensors were expressed using the previously described pre-pro-glucagon specific Perceval expression construct as expression backbone (14). Cloning was performed using homologous recombination in E. coli. The whole vector backbone including pre-pro-glucagon (ppGCG) specific UTRs, promoter and terminator sequence was amplified via PCR (Table 1). In separate PCRs, the insert sequences of Grx1-RhoGFP (22) and mORP (22) were amplified using primers with overhangs complementary to the ppGCG specific UTR sequences of the Vector backbone. The linear insert with complementary overhangs as well as the linear vector backbone was purified from a 1% Agarose gel via electrophoresis (Macherey-Nagel #740609.50) and heat-shock transformed in OmniMAX *E. Coli*. Bacteria were grown for 6h in SOC media to allow homologous recombination before plating on kanamycin plates. Liquid stocks were inoculated from overnight colonies, plasmid was isolated on mini prep scale (Macherey-Nagel #740588.50) and sequenced before sending the plasmid for viral packaging to Vector Biolabs (293 great Valley Parkway Malvern, PA 19355).

**Table 1.**
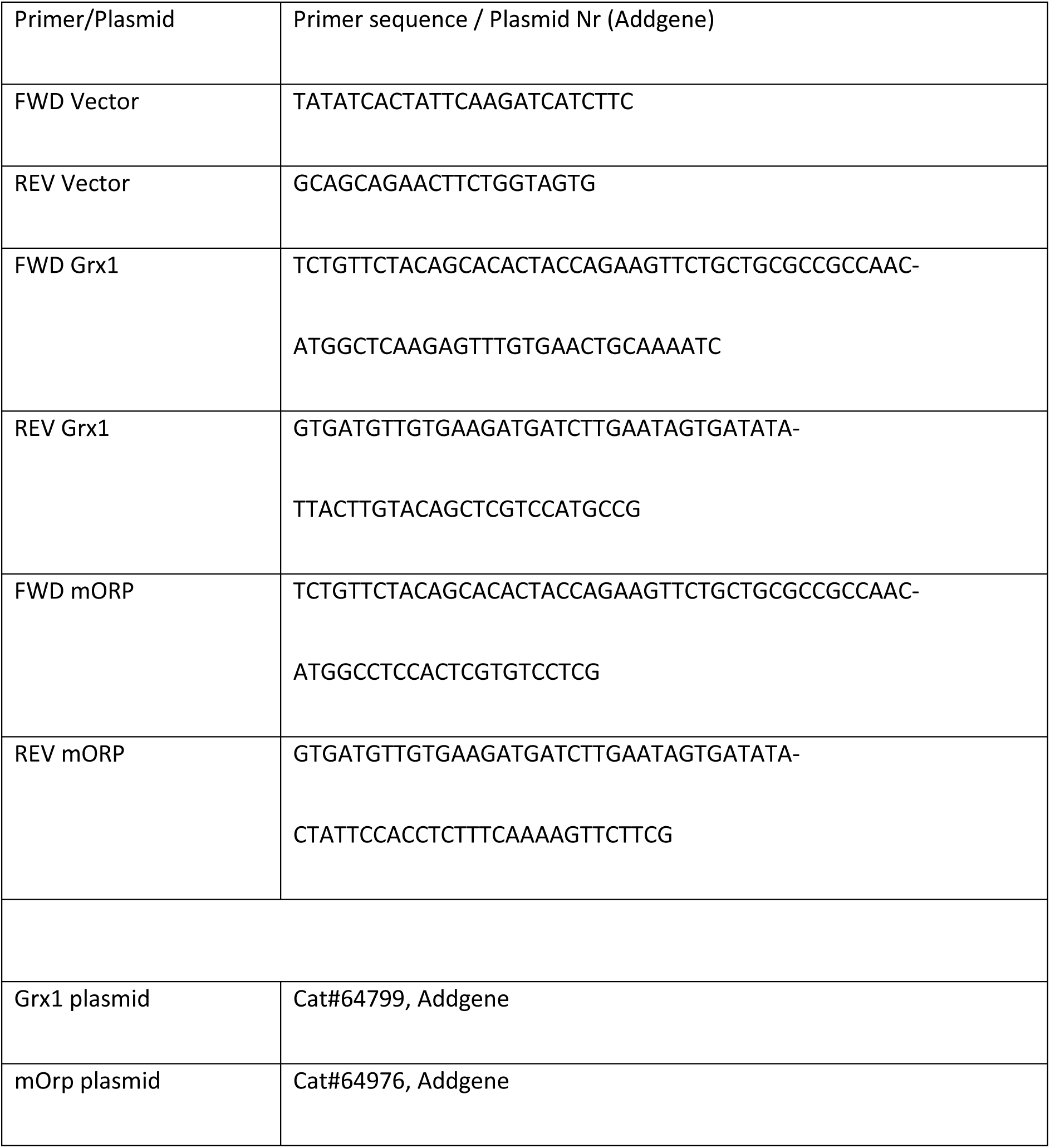
Primer sequences for cloning.

### Live cell imaging

Islets were washed in RPMI1640 media (Gibco, 11879-020) supplemented with 7mM glucose (without FBS or P/S). After washing, islets were transduced with recombinant adeno viruses for 2 hours in FBS-free RPMI1640 media. The transduction was then terminated by adding FBS containing media in a 1:1 ratio. Islets were incubated overnight and transferred to fresh media the following day. Before imaging, islets were transferred to KRB supplemented with a physiological fatty acid mixture as previously described (14) and incubated at 37°C incubator with 5% CO_2_ for 1h. A Leica SP5-X with a 40× objective, a 405 nm pulsed diode laser, white light laser, HyD detector and PMT detectors were used. Perceval, Grx and mORP were all detected using Leica standard GFP settings. Grx-roGFP and mOrp-roGFP probes were excited at 405 nm and 488 nm in a sequential setup and Perceval at 500 nm. Images were collected as xyzt stacks every 2.5 minutes with 2.5 µM z intervals. Changes of glucose concentration or addition of pharmacological compounds are indicated in the relevant figure legends.

PKA imaging was performed by virally transducing whole islets with an AKAR4 sensor cloned into the above described ppGCG cassette (23). Islets were picked into KRB and left to pre-incubate at 37°C, 5% CO_2_ for 30 min prior to imaging as indicated in figures and figure legends. Images were collected using a Nikon Ti2 CREST V3 spinning disk with a 40x objective as a xyzt stack every 2.5 min, excited at 446 nm and emission collected at 471 nm and 539 nm. Islets were perifused with KRB containing 1 mM glucose.

Calcium imaging was performed by loading islets with Fluo-4 (4 µM) (Thermo Fisher Scientific, F14201) and pluronic acid (2.5 µl per ml) in KRB at room temperature for 30 min (Leica SP5-X) or 70 min at 37°C, 5% CO_2_ (Nikon Tie2 CREST V3 spinning disc). Images were collected at a sampling rate of 2 images per second, excited at either 488 nm or 476 nm and emission was collected at 510 nm. Islets were pre-incubated and subsequently perifused with KRB buffer as indicated in figures and figure legends. Calcium imaging data were analysed in MATLAB using the findpeaks() function of the signal processing toolbox.

### Hormone secretion assays

Islets were size matched and picked into 100 µl KRB containing glucose and additives N-acetyl-cysteine (NAC) (Sigma Aldrich A7250), diamide (Sigma Aldrich, D3648), dehydroepiandrosterone (DHEA) (Sigma Aldrich, 700087P), as indicated and pre-incubated for 1h at 37°C, 5% CO_2_. Unless stated otherwise, (anti)oxidants and inhibitors were applied throughout the experiment. After pre-incubation, the supernatant was removed and islets washed once in 1mM glucose KRB. Following this, 100 µl 1 mM glucose KRB were added and islets were incubated for 1h at 37°C, 5% CO_2_. The supernatant was subsequently collected and stored at −80°C and islets washed once in 5 mM glucose KRB. Following this, 100 µl 5 mM glucose KRB was added and islets were incubated for 1h at 37°C, 5% CO_2_. The supernatant was then again removed and stored at −80°C, while the islets were lysed by sonication in acidified ethanol and stored in the same solution at −80°C for later hormone content measurements. Insulin and glucagon were measured using MSD (Meso Scale Discovery) or HTRF (Revvity) assays according to the manufacturer’s instructions. Somatostatin was measured using radio-immuno-assay as described previously (24).

### In vivo experiments

To explore NAC feeding in lean mice, C57B6NRj female mice on regular chow diet were provided with either water or water containing NAC (2 mg/ml) (n=8). The water was changed three times per week for 6 weeks; no changes in water consumption was observed. To explore the effects of NAC during high fat diet feeding, male C57B6NRj mice were fed a either a control (#D12450J, Research diets) or a high fat diet with 60% of calories derived from fat (#D12492, Research diets) (n=4) for six weeks and were provided with either water or water containing NAC (2 mg/ml) as described above.

### Tolerance tests

To test glucose tolerance (GTT), mice were fasted for 6h prior to the glucose injection (2 g/kg). For 2-deoxyglucose (2-DG) and insulin tolerance tests (ITT), the fasting period was 4 h prior to injecting 2-DG (500 mg/kg) or Insulin (0.75 IU/kg; Actrapid, Novo). All injection solutions were prepared in PBS and sterile filtered. During GTT, ITT and 2-DG tests, blood for glucose and hormone measurements were obtained right before and 30 min after injection. Blood glucose was monitored in 30 min intervals throughout (Contour BGMs, Bayer). Blood was collected on ice in tubes containing aprotinin (A1153, Sigma Aldrich), and plasma was prepared by centrifugation at 2600g at 4°C for 15 min and stored at −80°C.

### Fatty acid oxidation

Fatty acid oxidation was determined as previously described (14) using radiolabelled palmitate (NET043001MC; Perkin Elmer).

### Cell culture and immunoblotting

Alpha TC-1 6 cells (ATCC #CRL-2934) were cultured in RPMI 1640 media (Gibco,11879-020), with 10 mM Glucose (Sigma, G8769), 15 mM HEPES (Gibco,15630-056), 10% FBS (Gibco, 10270-106) and 1% P/S (Gibco,15140-122) at 5% CO_2_ and 37°C. The cells were cultured in 6 well plates for 72 h and then washed twice with KRB buffer containing 1 mM glucose. Cells were incubated in KRB buffer supplemented with NAC (300 µM) and Forskolin (1 µM) for 1 h at 37°C and 5% CO_2_. Cells were lysed in lysis buffer (150 mM NaCl, 20 mM Hepes, 1 mM EDTA, 10% glycerol, 0.5% Triton X-100, 1% SDS, protease and phosphatase inhibitor cocktail (Thermo Fisher Scientific, Cat. No. 78440)), sonicated 3 times for 10 sec each, and centrifuged at 15,000 g for 15 min. Cell lysates were collected and stored at −80°C. Protein concentrations were measured using a Bradford Assay (Bio-Rad, DC protein assay Reagent B, # 5000114 DC Protein Assay Reagent A # 5000113, DC Protein Assay Reagent S # 5000115). Samples were prepared by adding 50 mM dithiothreitol (DTT), sample buffer (NuPAGE, Cat. No. NP0007, Invitrogen) and heating at 95°C for 5 minutes. Protein samples (20 µg) were loaded in polyacrylamide gels (NuPAGE 4-10% Bis-Tris 12 well, Thermo Fisher Scientific Cat. No. NP0322box) and then transferred to a PVDF membrane (Invitrogen, Thermo Fisher Scientific Cat. No. LC2005). Membranes were stained with Ponceau stain and later blocked in Tris-buffered saline with 0.1% Tween 20 (TBST) buffer with 5% skimmed milk for 1 h at RT on a shaker. Primary antibodies (Table 2) were added overnight at 4°C. The next day, membranes were washed three times with TBST for 5 min each, followed by incubation with appropriate HRP conjugated secondary antibodies. Bands were visualized with Fusion FX Spectra (Vilber Lourmat, Eberhardzell, DE).

**Table 2.**
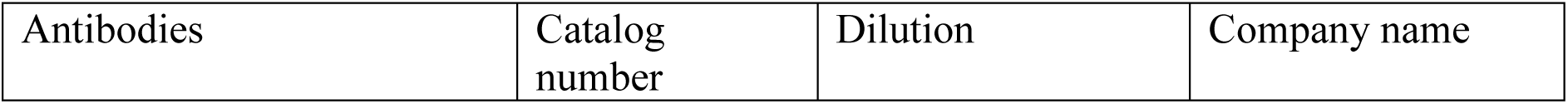
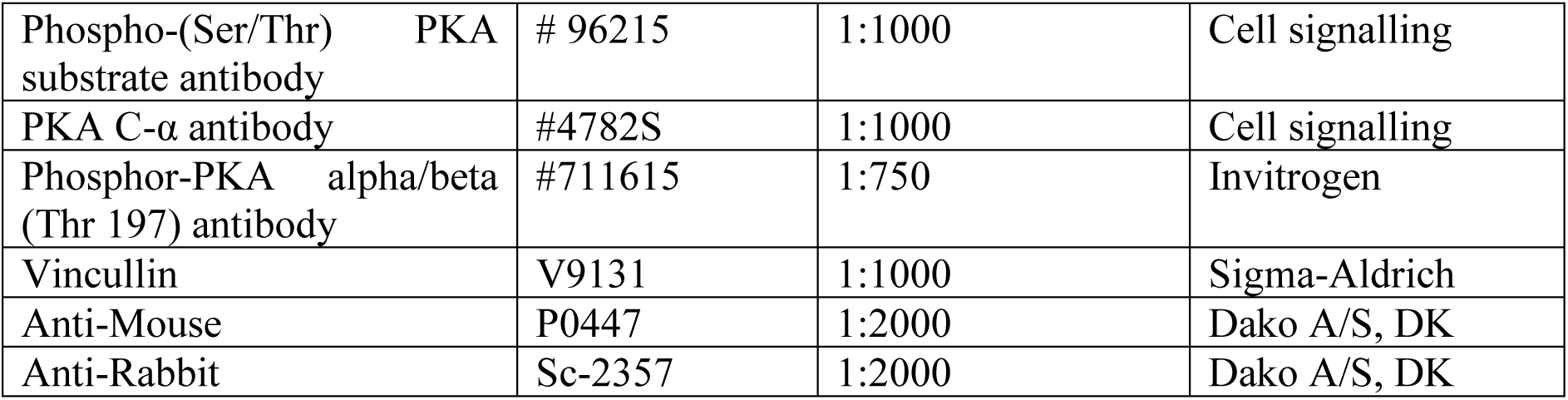
Antibodies used for western blotting.

### Statistical analysis

Statistical analysis was performed in GraphPad Prism 10. A Students t-test was used to compare two groups, a one-way ANOVA to test differences between more than two groups and a two-way ANOVA to compare groups characterized by two variables. Data were tested for equal variance and normality. When either assumption was not met, data were log-transformed and plotted as geometric means. Outliers were removed using ±2SD as the cut-off.

## Results

### Increased extracellular glucose elevates the redox potential to maintain glucagon release

Cytosolic glucose metabolism can be divided into three main pathways, glycolysis, gluconeogenesis and the pentose phosphate pathway (PPP). In glycolysis, glucose is converted to pyruvate which is then either oxidised in the mitochondria or converted to lactate. The PPP provides metabolites for nucleotide synthesis but is also the main provider of reductive equivalents in the form of NADPH. These reductive equivalents can be used either for biosynthesis, such as fatty acid synthesis (FAS), or for reducing oxidized glutathione (GSSG) to maintain intracellular redox potential (Fig. 1A). While FAS is required to maintain alpha cell size and function (25), it is unknown whether changes in GSSG play a role in the regulation of glucagon secretion. To understand whether glucose is metabolised through the PPP in alpha cells, we incubated αTC1-6 cells in 1 or 16.7 mM glucose for 1h and measured ribose-5-phosphate (Rib5P) using GC/MS (26). Under these conditions the elevation in glucose led to a robust increase in ribose-5-phosphate content (Fig. 1B), in line with previous observations in whole rat islets (27). We therefore hypothesised that increases in extracellular glucose would lead to elevations in reduced glutathione (GSH) and, thus, cytosolic redox potential in alpha cells. To explore this, we transduced primary isolated islets with an adenovirus expressing the glutathione redox sensor Grx1- roGFP under the control of the pre-pro-glucagon promoter (supplementary table 1) (14; 22). Time lapse images were captured from transduced islets perifused with increasing glucose concentrations. The gradual elevation in glucose from 1 to 5 to 20 mM glucose led to an acute increase in cytosolic redox potential (Fig. 1B-D). This suggested that elevations in glucose may to acutely increase the cytosolic redox potential through the PPP. We therefore tested the effect of the glucose 6 phosphate dehydrogenase (G6PDH) inhibitor, dehydroepiandrosterone (DHEA) (27), on glucagon secretion. Treatment with DHEA led to a reduction in glucagon secretion at 1 mM glucose but did not affect glucose-induced inhibition of glucagon secretion (Fig. 1F). While the more specific and potent G6PDH inhibitor G6PDi (28), lowered glucagon secretion at 1mM glucose and 5mM glucose, removing the inhibitory effect of glucose on glucagon secretion (Fig 1G). Together with previous observations that DHEA can block glucose-induced increases in Rib5P and GSH in rat islets (27), this supports that metabolism of glucose through the PPP in alpha cells is important for maintaining glucagon secretion at low glucose. In line with this, when the cytosolic redox potential was depleted by pre-incubation of whole islets in 1 mM glucose, glucagon release lowered at low glucose (1 mM) (Fig. 1H and I). This did not affect insulin secretion (Supplementary Fig. S1B).

**Figure 1:**
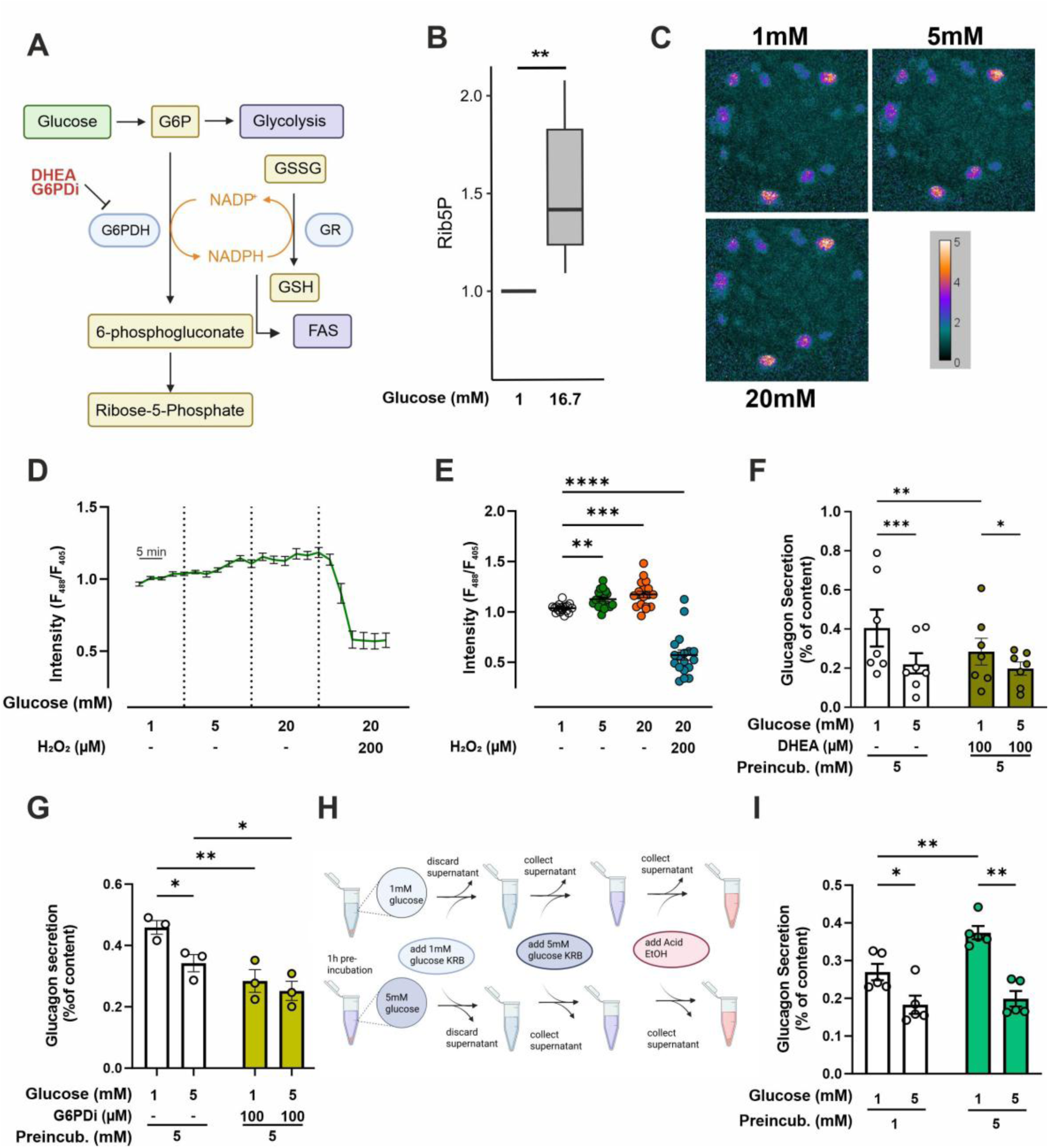
Increased extracellular glucose elevates the redox potential in alpha cells; (A) Schematic of cytosolic glucose metabolism. (B) Ribose 5 phosphate (Rib5P) measured in αTC1-6 cells after incubation in 1 and 16 mM glucose (n=18 cells from 3 mice in 3 independent experiments). (C) Representative ratio-metric heatmap images and (D) average traces of whole pancreatic islets infected with adenoviral constructs expressing the redox sensor Grx1-roGFP under the GCG promoter perifused with 1, 5 and 20 mM glucose (n=18 cells from 3 mice in 3 independent experiments). (E) Accumulated data from D. (F) Glucagon secretion from whole islets pre-incubated in 5 mM glucose measured in response to 1 and 5 mM glucose with or without 100 µM DHEA (n=7 mice). (G) Glucagon secretion from whole islets pre-incubated in 5 mM glucose measured in response to 1 and 5 mM glucose with or without 100 µM G6PDi (n=3 mice). (H) Schematic of (I) Glucagon secretion from whole islets pre-incubated in 1 or 5 mM glucose measured in response to 1 and 5 mM (n=5 mice). All Data are presented as Mean ± SEM, * (P<0.05), ** (P<0.01), *** (P<0.001), **** (P<0.0001).

We next tested whether changes in cytosolic redox potential alone could affect glucagon secretion at low glucose. Pre-incubation of whole islets in 1 mM glucose with 300 µM of the antioxidant NAC (29) elevated the redox potential in the cytosol (Fig. 2A and B), similar to the increase observed with 5 mM glucose (Fig. 2A and B), suggesting that the addition of 300 µM NAC produced a physiologically relevant change in the redox potential. Pre-incubation in 1 mM glucose with 300 µM NAC also increased glucagon secretion at low glucose (Fig. 2C) without changing insulin or somatostatin release (Supplementary Fig. S2A and B). In line with these observations, applying hydrogen peroxide (H_2_O_2_) led to a concentration-dependent stepwise decrease of the redox potential in alpha cells (Fig. 2G and H) and the addition of low concentrations (25 µM) of H_2_O_2_ during secretion lowered glucagon release at low glucose in islets pre-incubated in 5 mM glucose (Fig. 2I). These findings indicate that, in mouse alpha cells, glucose is metabolised through the PPP, which leads to an increase in the cytosolic redox potential, and amplification of glucagon secretion at low glucose.

**Figure 2.**
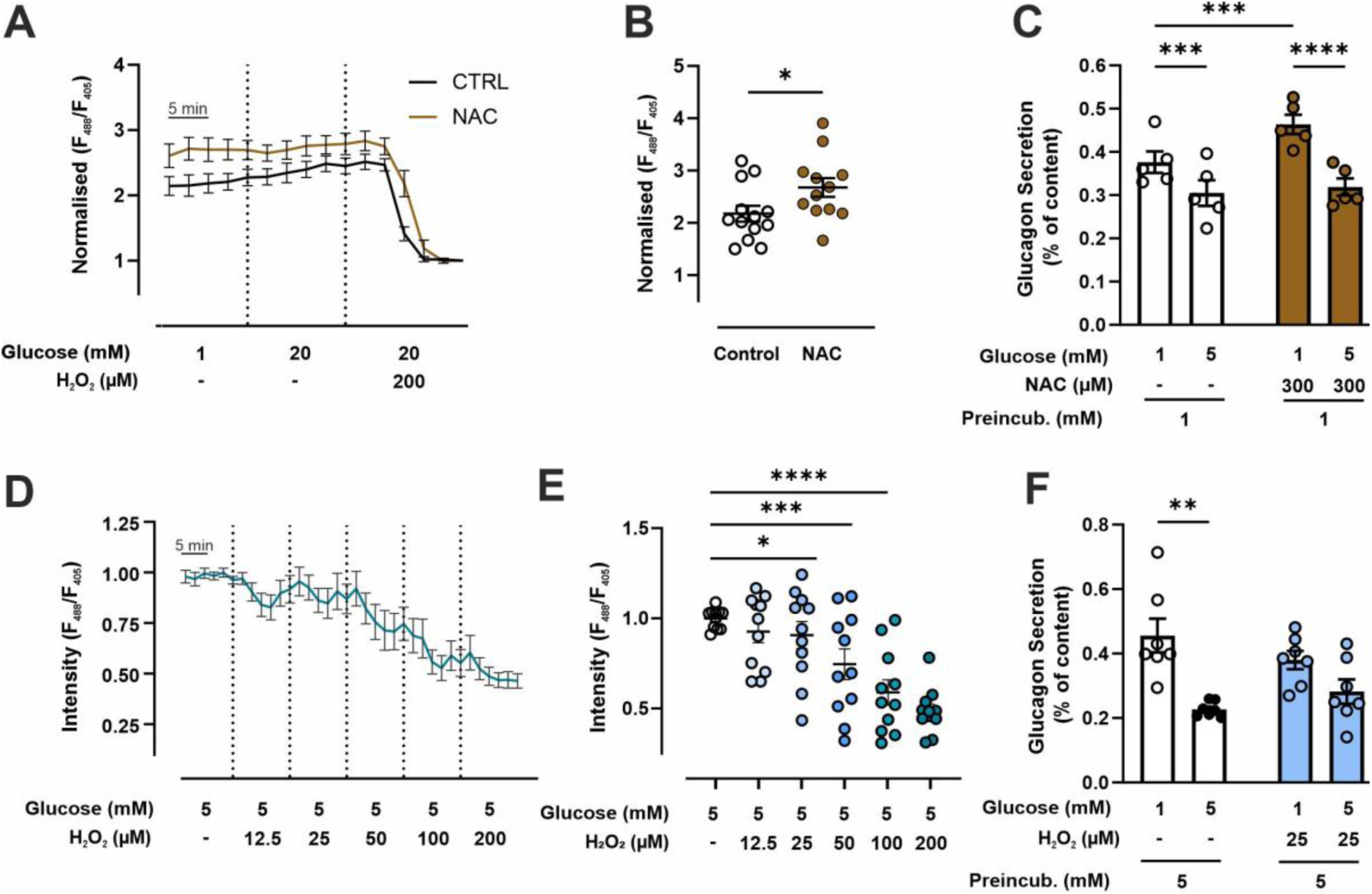
Glucagon secretion at low glucose depends on the cytosolic redox potential. (A) Live cell imaging of redox potential in alpha cells in whole islets infected with PPGCG-promoter-Grx1-roGFP, pre-incubated for one hour in 1 mM glucose with or without 300 µM NAC and perifused with 1, 5 or 20 mM glucose with or without 300 µM NAC (n= 12 cells from 3 mice in 3 independent experiments). (B) Accumulated data from A. (C) Glucagon secretion in whole islets, pre-incubated in 1 mM glucose with or without 300 µM NAC, measured in response to 1 and 5 mM glucose with or without 300 µM NAC (n=5 mice). (D) Live cell imaging of redox potential in alpha cells in whole islets infected with PPGCG-promoter-Grx1-roGFP, pre-incubated for one hour in 5 mM glucose and perifused with 5 mM glucose with increasing concentrations of H_2_O_2_ (12.5-200 µM) (n= 11 cells from 3 mice in 3 independent experiments). (E) Accumulated data from D. (F) Glucagon secretion in whole islets, pre-incubated in 1 mM glucose, measured in response to 1 and 5 mM glucose with or without 25 µM H_2_O_2_ (n=5 mice). All Data are presented as Mean ± SEM, * (P<0.05), ** (P<0.01), *** (P<0.001) **** (P<0.001).

### Changes in glucose do not affect mitochondrial H_2_O_2_ production

We next wanted to understand how changes in the cytosolic redox potential could affect glucagon secretion. Alpha cells rely on fatty acid oxidation to maintain intracellular ATP and they are therefore subject to a higher production of reactive oxygen species (ROS) in the electron transport chain (30). Interestingly, ROS arising from the electron transport chain has been suggested to lower exocytosis in alpha cells (31), We therefore speculated that the increased redox potential in the cytosol may alleviate production of ROS in the mitochondria. To explore whether mitochondrial production of reactive oxygen species was affected by changes in cytosolic redox potential, we used the mitochondrial H_2_O_2_ probe mOrp-roGFP under the control of the pre-pro-glucagon promoter (Supplementary table 1) (14; 32). We first tested whether the probe could detect changes in mitochondrial H_2_O_2_ in response to extracellular increases in H_2_O_2_. In whole islets, application of 50 µM H_2_O_2_ was needed to increase mitochondrial H_2_O_2_ (Fig. 3A). Elevating glucose from 1 to 20 mM in whole islets pre-incubated in 1 mM glucose had no impact on mitochondrial H_2_O_2_ levels in alpha cells, neither did pre-incubation with 300 µM NAC (Fig. 3B), suggesting that changes in glucose does not affect the mitochondrial redox potential in alpha cells. In line with previous findings, this suggests that alpha cells express high levels of antioxidant enzymes such as glutathione peroxidase and catalase (33). Despite this, the elevated secretion observed with 300 µM NAC and 5 mM glucose pre-incubation could still be driven by changes in substrate oxidation. Pre-incubation in NAC at 1 mM glucose resulted in an increased ATP/ADP ratio (Fig. 3C-D) measured using an alpha cell specific version of the PercevalHR probe (14), without changes in fatty acid oxidation at low glucose (Fig. 3E). This indicates that the redox driven effect of NAC or glucose are independent of mitochondrial redox state but contributes to increase the ATP/ADP ratio in the cytosol at low glucose to support glucagon secretion.

**Figure 3:**
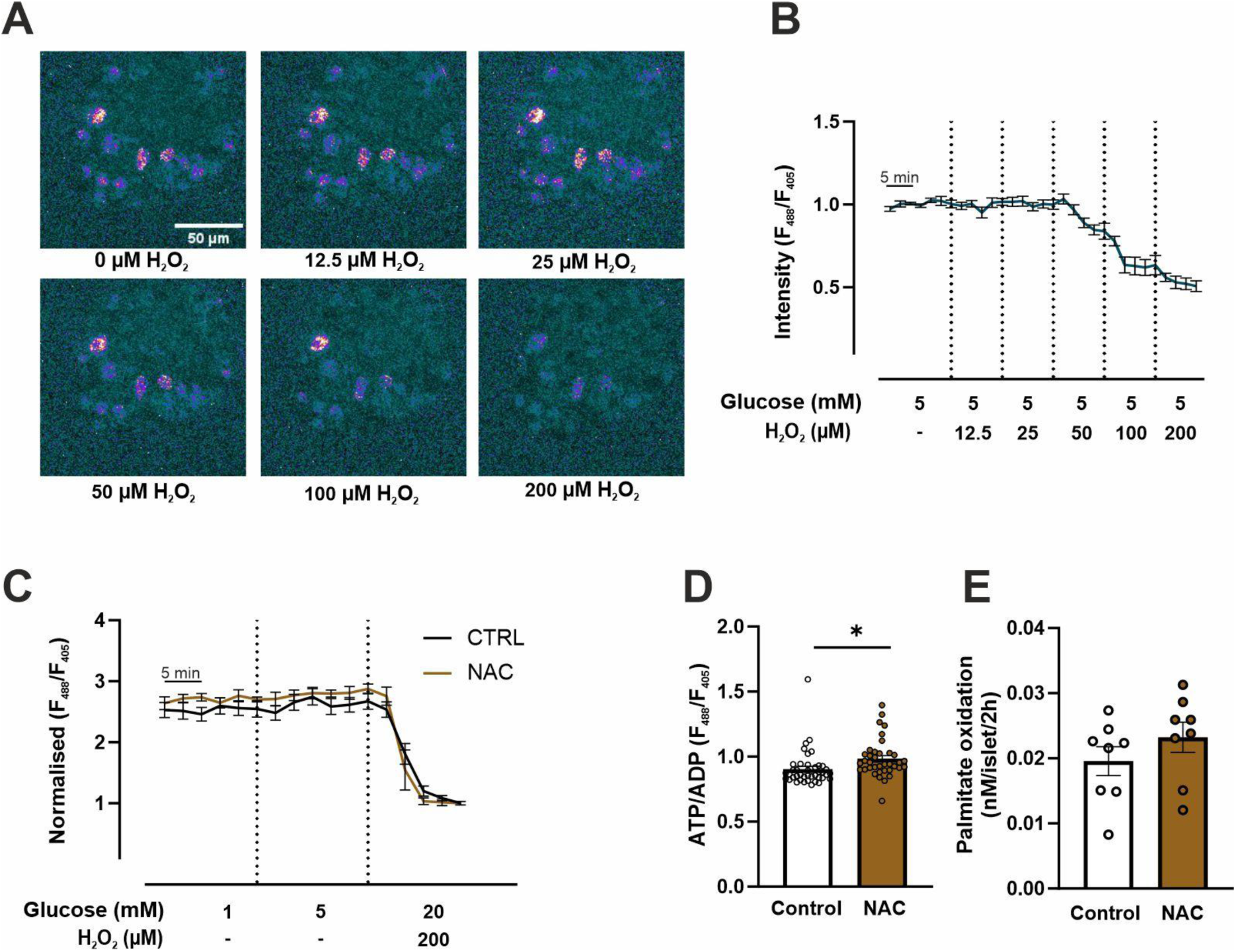
Higher extracellular glucose does not affect mitochondrial H_2_O_2_ production. (A) representative images from (B) Live cell imaging of mitochondrial H_2_O_2_ in alpha cells in whole islets, infected with PPGCG promoter-mOrp1-roGFP and pre-incubated for one hour in 5 mM glucose and perifused with 5 mM glucose with increasing concentrations of H_2_O_2_ (12.5-200 µM) (n=8 cells from 2 mice in 3 independent experiments). (B) Live cell imaging of Mitochondrial H_2_O_2_ in alpha cells in whole islets, infected with PPGCG promoter-mOrp1-roGFP and pre-incubated for one hour in 1mM glucose with or without 300 µM NAC and perifused with 1 or 5 mM glucose (n=13-14 cells from 3 mice in 3 independent experiments). (C) Accumulated data from live cell imaging of on ATP/ADP in alpha cells in whole islets, infected with PPGCG promoter-PercevalHR and pre-incubated for one hour in 1mM glucose or with 1 mM glucose and 300 µM NAC (n= 40 cells from 2 mice in 3 independent experiments). (D) Beta oxidation measured using 9,10^3^H-Palmitate in whole islets pre-incubated in 1 mM glucose with or without 300 µM NAC (n=8 mice). All Data are presented as Mean ± SEM, * (P<0.05).

### Changes in the redox potential regulate protein kinase A activity in alpha cells

Roughly, 10 % of all kinases have been hypothesised to be regulated by changes in redox potential (34). In protein kinase A (PKA), oxidation of two free cysteines (Cys199 and Cys343) in the catalytic subunit (35; 36) leads to dephosphorylation of Thr179 (36) which is critical for PKA activation (Fig. 4A) (37). PKA plays a key role in mediating the effects of adrenaline, glucose, and somatostatin on glucagon release (10; 12; 16; 38; 39). We therefore hypothesised that the effects of changes in cytosolic redox potential on glucagon secretion could be driven by changes in PKA activity. To test this, we first challenged alpha cells with diamide, a sulfhydryl specific oxidant targeting all free cysteine residues. Incubating mouse islets with diamide (10 μM) abolished the potentiating effect of 5 mM glucose pre-incubation on glucagon secretion (Fig. 4B). This aligns with previous findings showing that diamide and modest increases in oxidative state can decrease PKA activity (36). Incubating αTC1-6 cells in 1 mM glucose with 300 µM NAC resulted in increased phosphorylation of Thr179 (Fig. 4C) and led to a 40% increase in PKA substrate phosphorylation compared to cells incubated in 1 mM glucose without NAC (Fig. 4D). We then tested whether PKA activity in alpha cells in whole islets was altered in response to pre-incubation in 5 mM glucose. Using an alpha cell specific version of the PKA activity probe AKAR4 (Supplementary Table 1A) (23), we captured time lapse images of whole islets every 2.5 min for 10 min immediately after placing islets in 1 mM glucose. Under these conditions, PKA activity was increased in alpha cells in islets pre-incubated in 5 mM glucose compared to alpha cells in islets pre-incubated in 1 mM glucose (Fig. 4E and F). This suggested that the elevated redox potential induced by pre-incubation in 5 mM glucose led to increased PKA activation. In line with this, the effect of 5 mM glucose pre-incubation on glucagon secretion was abolished by the PKA inhibitor H89 (Fig. 4G). These findings suggest that during euglycemia, glucose metabolism loads alpha cells with redox equivalents, and that this reduced environment maintains PKA activity and glucagon secretion during subsequent exposure to hypoglycaemia.

**Figure 4:**
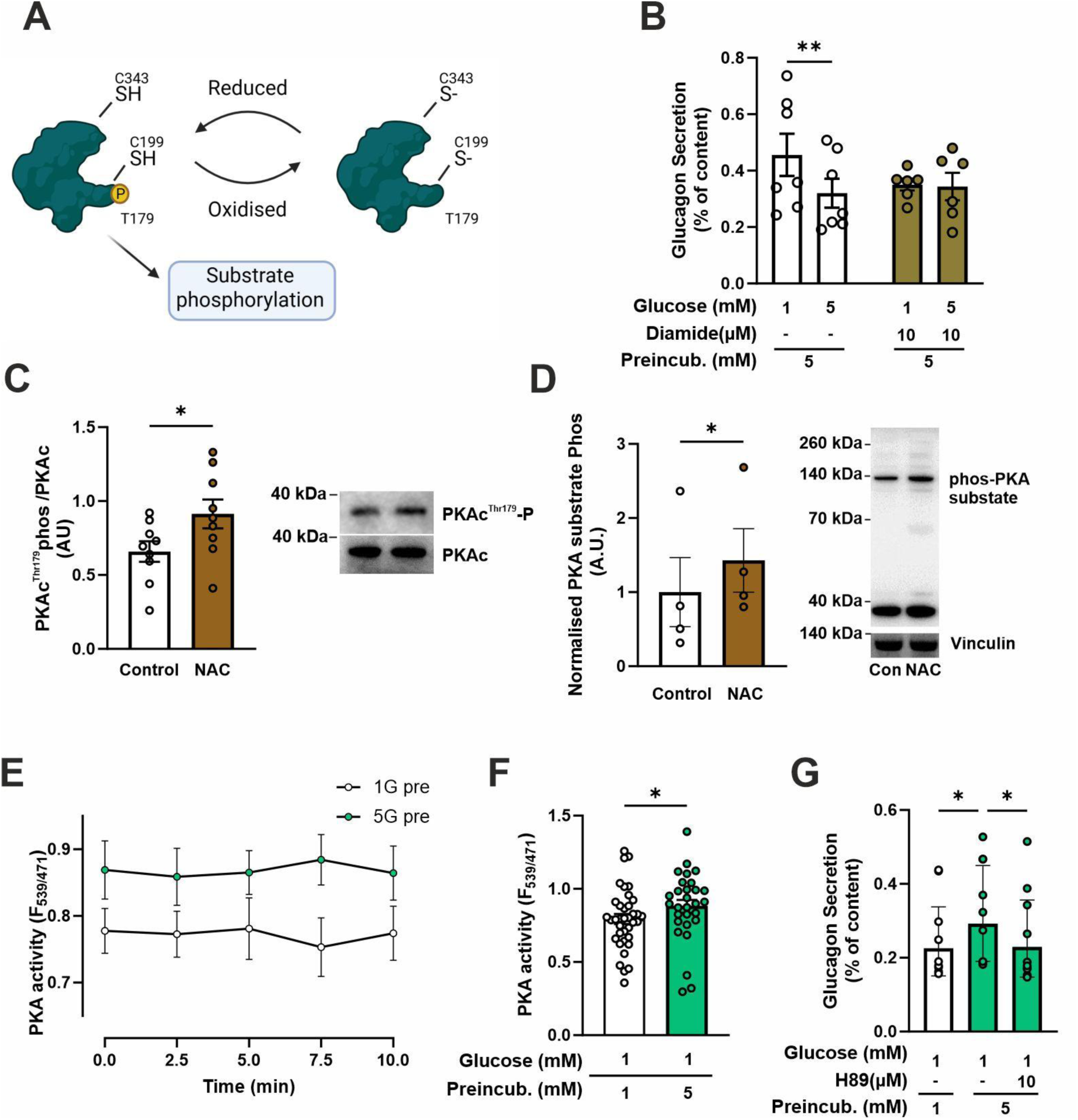
Changes in the redox potential regulate Protein kinase A activity in alpha cells. (A) schematic of how oxidation regulates of PKAc activity. (B) Glucagon secretion in whole islets pre-incubated in 1 mM glucose measured in response to 1 and 5 mM glucose with or without 10 µM Diamide (n=7 mice). (C) Quantification of PKAc^thr179^ phosphorylation in alphaTC1-6 cells in response incubation in 1 mM glucose with or without NAC (n=9 independent experiments). (D) Quantification of PKA substrate phosphorylation in alphaTC1-6 cells in response incubation in 1mM glucose with or without 300 µM NAC (N=3 independent experiments). (E) Average PKA activity in alphaTC1-6 cells at 1mM glucose from whole islets pre-incubated in 1 or 5 mM glucose (n=30-38 cells from 4 mice in 4 independent experiments). (F) Average data from (E). (G) Glucagon secretion from whole islets pre-incubated in 1 or 5 mM glucose with or without 10 µM H89 (n= 10 mice). All Data are presented as Mean ± SEM except (G) where data are represented as geometric mean ± SD, * (P<0.05), ** (P<0.01), **** (P<0.0001).

### Increases in the cytosol redox potential activates alpha cells at low glucose

Both PKA (38; 40) and ROS (41) have been shown to activate L-type calcium channels. We e therefore explored the effects of the increases in the cytosolic redox potential on intracellular calcium responses to glucose in alpha cells. Using activity in 1 mM glucose to identify putative alpha cells, we measured calcium oscillations in control and NAC treated whole islets. In control islets, the calcium oscillation frequency was reduced in 5 mM glucose (Fig. 5A). Treatment of whole islets with NAC did not affect the calcium oscillation frequency in 1 mM glucose, but increasing glucose concentration to 5 mM failed to reduce the average oscillation frequency (Fig. 5A). We therefore wondered whether NAC treatment led to activation of other cell types, such as beta or delta cells, that are not normally active at 1 mM glucose. However, NAC had no effect on insulin or somatostatin release at low glucose (Supplemental Fig. 2A and B). The capacity of glucose to increase calcium oscillations in alpha cells is a known phenomenon (42; 43). We therefore speculated that the absence of a glucose effect was due to a subpopulation of cells which were insensitive to 5 mM glucose, something that would be revealed by exploring the median, rather than the mean, calcium oscillation frequency. In both control and NAC-treated islets, glucose lowered the median calcium oscillation frequency (Fig. 5B), with most cells showing a reduction in calcium oscillation frequency in response to elevated glucose, while a subset of cells did not respond (Fig. 5C). Analyses of the two subpopulations separately revealed two distinct populations with differential responses to the increase in glucose from 1 to 5 mM, with one population clearly lowering peak frequency and one maintaining oscillatory activity throughout the recording (Fig. 5D-E). Regardless, NAC did not affect average peak frequency during 1 mM glucose (Fig. 5D). However, when analysing the number of active cells, there were 3 times as many active cells in NAC treated islets compared with control islets in 1 mM glucose (Fig. 5F). Calcium oscillations in both control and NAC treated islets could be blocked by the addition of the L-type calcium channel inhibitor isradipine (Fig. 5G), which also lowered glucagon secretion from islets pre-incubated in 5 mM glucose (Fig. 5H). These findings are in line with previous observations that H_2_O_2_ lowers and GSH increases calcium influx in alpha cells (31) and suggest that redox-mediated amplification of glucagon secretion depends on L-type calcium channel activity.

**Figure 5.**
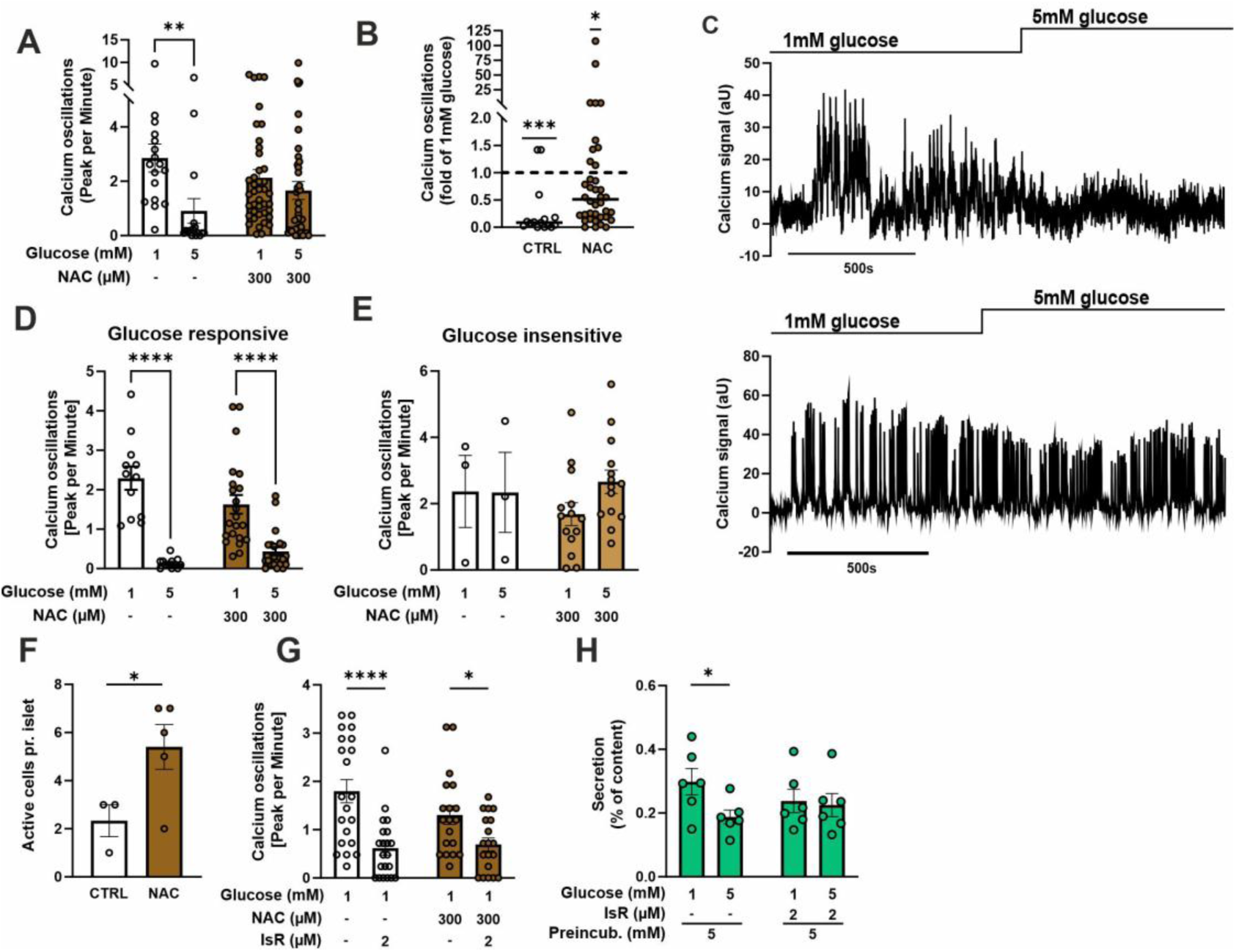
A more reduced cytosol activates alpha cells at low glucose. (A) Calcium peak frequency of putative alpha cells (active in 1mM glucose) in islets pre-incubated for one hour in 1 mM glucose with or without 300 µM NAC and perifused with 1 or 5 mM glucose in 1 mM glucose with or without 300 µM NAC (n= 15-41 cells from 3 mice in 3 independent experiments). (B) Median calcium peak frequency fold change from 1 to 5 mM glucose with or without 300 µM NAC in putative alpha cells in islets pre-incubated for one hour in 1 mM glucose with or without 300 µM NAC (Data from (A)). (C) Representative traces of intracellular calcium in glucose responsive cells (top) and glucose insensitive cells (bottom). (D and E) Calcium peak frequency of glucose responsive and insensitive cells from (B). (F) Average number of active cells pr islet at 1mM glucose in islets pre-incubated for one hour in 1 mM glucose with or without 300 µM NAC (n=3-5 islets from 3 mice). (G) Calcium peak frequency of putative alpha cells in islets pre-incubated for one hour in 1 mM glucose with or without 300 µM NAC and perifused with 1 mM glucose in 1 mM glucose with or without 300 µM NAC and 2 µM Isradipine (n= 19-21 cells from 3 mice in 3 independent experiments). (H) Glucagon secretion islets pre-incubated for one hour in 5 mM glucose at 1 and 5 mM glucose with or without 2 µM Isradipine (n=6 mice). All Data are presented as Mean ± SEM, * (P<0.05), ** (P<0.01), **** (P<0.0001).

### A reduced environment affects glucagon levels in vivo

To understand whether the increased redox potential and subsequent glucagon secretion observed in the islets impact on circulating glucagon levels, we supplemented healthy chow fed female C57B6Nrj mice with NAC (2 mg/ml) in the drinking water for 6 weeks. The addition of NAC to the drinking water did not affect body weight or body fat percentage (Fig. 6A and B). Despite this, a perceived hypoglycaemia test (injection with 2-deoxy glucose (2DG) 50mg/kg) stimulated glucagon secretion 7.5-fold in NAC treated animals compared to 4.5-fold in controls (Fig. 6 C-E). This suggested that NAC treatment could alter plasma glucagon dynamics. To understand whether this could affect whole body glucose metabolism, NAC fed animals were subjected to an intraperitoneal glucose tolerance test (IPGTT). The glucose excursion caused by injecting 2 mg of glucose/kg body weight was similar in control and NAC treated animals, but NAC fed mice returned to baseline glucose levels faster than control mice (Fig. 6 F and G); an effect which was not caused by changes in insulin secretion (Fig. 2H). While control mice showed a slight increase in plasma glucagon 30min after the glucose bolus, NAC treated animals slightly lowered plasma glucagon 30 min after glucose (Fig. 2J and K). Combined with the findings from isolated islets, this suggests that a reduced environment in the cytosol of alpha cells leads to improved regulation of glucagon and possibly faster recovery from a glucose challenge.

**Figure 6.**
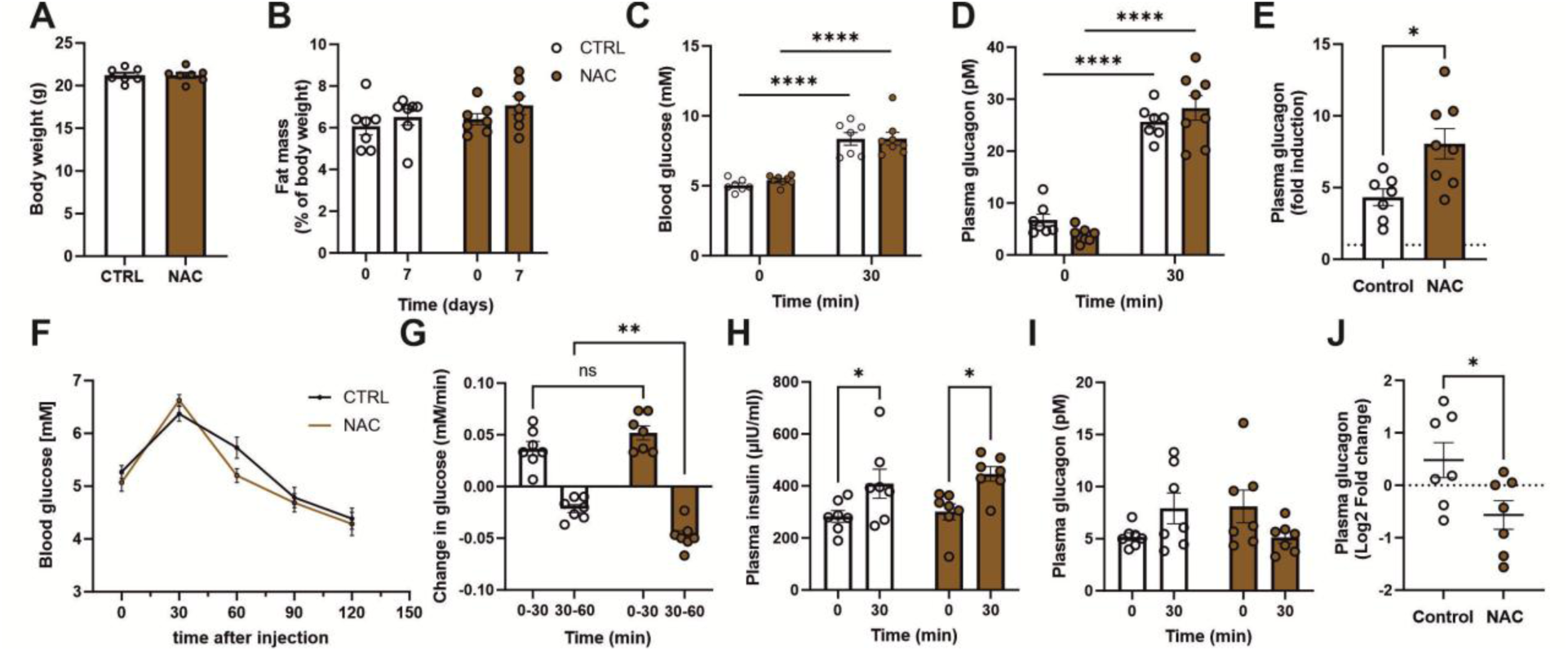
A reduced environment affects glucagon regulation *in vivo*. (A) body weight and (B) fat mass in C57B6Nrj female mice fed NAC (2mg/ml) in the drinking water for 6 weeks (n=7-8 Mice). (C) Blood glucose and (D) plasma glucagon levels in response to a 2-deoxy glucose injection after 6 weeks of NAC feeding (n=7 mice). (E) Fold change in glucagon from 0-30 min from D. (F-G) Blood glucose, (H) plasma insulin and (I) plasma glucagon levels in response to an IP glucose tolerance test after 7 weeks of NAC feeding (n=7 mice). (J) Fold change in glucagon from 0-30 min from (J). All Data are presented as Mean ± SEM, * (P<0.05), **(P<0.01), **** (P<0.0001).

## Discussion

Here we explored the role of non-oxidative glucose metabolism in alpha cell function. Our findings suggest that glucose in alpha cells is metabolised through the PPP to maintain the cytosolic redox potential under low glucose conditions. We also demonstrate that a more reduced potential is required for basal PKA activity and determines the magnitude of glucagon release in response to hypoglycaemia.

The role of glucose metabolism in the regulation of alpha cell function is widely debated (13; 14; 16; 44; 45). While previous studies have largely focused on the impact of glucose oxidation on ATP production (13; 46), other findings suggest that both glycolytic and gluconeogenic enzymes play important roles for the effects of glucose on glucagon secretion. Inhibiting or knocking out glucokinase, catalysing the initiating reaction of the glycolysis, leads to loss of glucose-inhibited glucagon secretion, while loss of the gluconeogenic gene G6Pase2 leads to increased inhibition by glucose (47; 48). Interestingly, the lower part of glycolysis has been suggested to be directly involved in the regulation of membrane potential through pyruvate kinase derived ATP production and closure of K_ATP_ channels (45) thereby lowering glucagon secretion. Rather than a direct regulation of glucagon secretion by glycolysis, we find that treatment with the G6PDH inhibitor DHEA leads to lower glucagon secretion at low glucose. These findings align with previous observations that, at higher glucose levels, alpha cells increase non-oxidative metabolism (17) and that loss of acetyl-CoA carboxylase 1 and FAS leads to impaired glucagon secretion, reduced glucagon content and cell size (25). Our findings demonstrate that glucose metabolism not only supports FAS (25) and regulation of ATP levels (13; 14; 19-21; 46; 49; 50), but play an important role in maintaining the cytosolic redox potential. In the present experiments, a more reduced redox potential induced by higher glucose enhanced glucagon secretion at low glucose. These results parallel findings in single human alpha cells where pre-incubation in 10 mM glucose amplifies glucagon secretion, while H_2_O_2_ applied via intracellular dialysis leads to impaired exocytosis (31). Rather than observing increased activity in single cells, we observed activity in a greater number of putative alpha cells, suggesting slightly different mechanisms in mouse and human alpha cells. Remarkably, both oscillation frequency and secretion could be inhibited by the addition of the L-type channel inhibitor isradipine. This contradicts previous findings where isradipine either had no effect (31) or stimulated glucagon release (51). There is no direct evidence to explain these differences, however, both fatty acids and changes in BSA can affect intracellular conditions (14; 52).

Both individuals with diabetes (31) and hyperglycaemic mouse models (49) show impaired glucagon release at low glucose. Previous findings suggest that this is due to impaired exocytosis, caused by increased production of ROS (31). The findings presented here suggest that the increased ROS could lead to a reduction in PKA activity, possibly contributing to the lower glucagon secretion observed at low glucose. Providing NAC to elevate reductive potential at the whole-body level also affected the fold-change of circulating glucagon levels, but not glucose handling. This is in line with observations in other models, showing marginal effects of changes circulating glucagon levels on circulating glucose levels (14; 19; 49). NAC treatment likely also affected the liver, brain and the sympathetic nervous system(53), all of which could contribute to the regulation of glucagon release and glucose handling.

In conclusion, we report here a so far unknown role for glucose as a potentiating factor for glucagon secretion. We find that under euglycemia, metabolism of glucose in the PPP elevates the cytosolic redox potential in alpha cells. This “loads” the cytosol with redox equivalents and plays an important role in maintaining PKA activity and glucagon secretion during hypoglycaemia.

## Author contributions

Conceptualization: JGK and AF, Data curation: AF and JGK, Formal analysis: AF, CLP, MHE, GD, HP, RI DBA, DN and GKP, Funding acquisition: JGK Investigation: AF, CLP, MHE, GD, HP, RI, DBA, DN, PS and GKP, Methodology: AF, PAP, and JGK, Resources: DAB, PS, JJH, PAP, AF and JGK, Supervision: PAP, JJH and JGK, Validation: AF and JGK, Visualization: AF and JGK, Writing Original Draft: AF and JGK, Writing – review and editing: JJH, PAP, PS, JGK, AF, CLP, MHE, GD, HP, RI, DBA, DN and GKP

## Data availability

All data required to assess our conclusions are included in the manuscript. Raw images and source data for all figures can be obtained upon reasonable request to the corresponding author (JGK).

## Acknowledgements

Imaging experiments were performed at the Centre for Advanced Bioimaging (CAB) at the University of Copenhagen.

---

**Supplementary figure 1:**
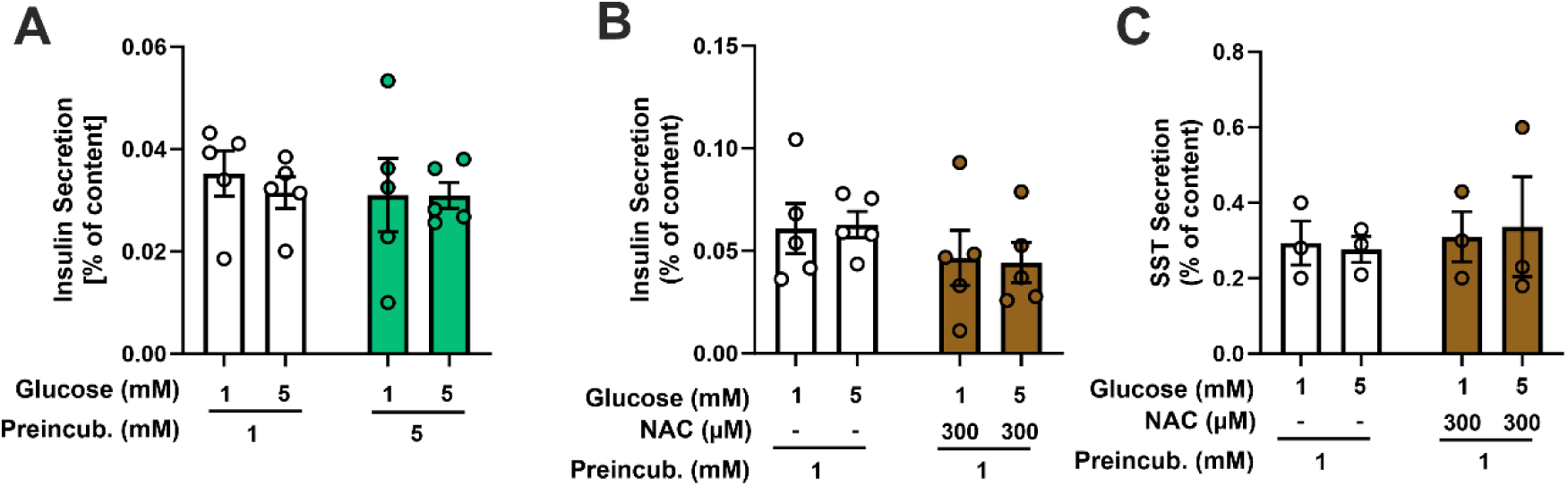
(A) Insulin secretion in response to 5 mM pre-incubation (n=5 mice). (B) Insulin and (C) somatostatin secretion in response to NAC treatment with 1 mM pre-incubation. All Data are presented as Mean ± SEM

**Supplementary Table 1:** Specificity of fluorescent probes used for live cell imaging calculated alpha cells as percent of cells expressing the probe (From 15-42 islets from 4 mice). Data are presented as Mean ± SEM.

| Probe | AKAR4 | mORP1 | Grx1 |
| --- | --- | --- | --- |
| Specificity | 95.7±1.4% | 92.4±3.6% | 100±0% |

